# Unified Framework for Representing and Ranking

**DOI:** 10.1101/086678

**Authors:** Jim Jing-Yan Wang, Halima Bensmail

**Author notes:** Corresponding author Email addresses (Jim Jing-Yan Wang), (Halima Bensmail).

## Abstract

In the database retrieval and nearest neighbor classification tasks, the two basic problems are to represent the query and database objects, and to learn the ranking scores of the database objects to the query. Many studies have been conducted for the representation learning and the ranking score learning problems, however, they are always learned independently from each other. In this paper, we argue that there are some inner relationships between the representation and ranking of database objects, and try to investigate their relationships by learning them in a unified way. To this end, we proposed the **U**nified framework for **R**epresentation and **R**anking (UR^2^) of objects for the database retrieval and nearest neighbor classification tasks. The learning of representation parameter and the ranking scores are modeled within one single unified objective function. The objective function is optimized alternately with regarding to representation parameter and the ranking scores. Based on the optimization results; iterative algorithms are developed to learn the representation parameter and the ranking scores on a unified way. Moreover, with two different formulas of representation (feature selection and subspace learning), we give two versions of UR2. The proposed algorithms are tested on two challenging tasks - MRI image based brain tumor retrieval and nearest neighbor classification based protein identification. The experiments show the advantage of the proposed unified framework over the state-of-the-art independent representation and ranking methods.

## 1. Introduction

In the database retrieval and nearest neighbor classification tasks, given a query object, we try to find some relevant objects from a database [1, 2]. The relevant objects here are defined as the objects of the same semantical class. For example, in the brain tumors diagnosis problem, given a tumor region in a Magnetic Resonance Imaging (MRI) image as a query, it could be very helpful for the diagnosis to retrieve tumors of the same pathological category from a brain MRI scans database [3]. While in drug discovery problem, given a query protein, it could also be useful to find the proteins sharing the same specific chemical properties or similar structure as the query protein from a protein database, so that they can be used as sources for the treatment [4]. To this end, in a typical database retrieval system, the feature vectors are usually first extracted from both the query and database objects, and then the query is compared against each database object to compute the similarities or dissimilarities using their feature vectors. Finally, all the database objects will be ranked according to their similarities to the queries in the descending order, and a few number of them with the largest similarities will be returned to the user, or used to make a classification decision. Because the similarity is used for ranking the database objects, it is also called ranking score [5].

Two fundamental problems have been studied widely is the learning of the representations of the objects feature vectors, and the learning of the ranking scores of the database objects to the query, as listed as follows:

- **Representation**: The original features extracted from the objects are usually very high-dimensional, redundant, sometimes noisy, and only occupying a part of the input space. Thus the original features may not capture the semantical information and could not be used directly to retrieve the relevant objects very well. In this case, it’s necessary to represent the feature vectors to another dataspace so that they could be represented better for the retrieval task. Many representation methods can be considered, such as feature selection [6], subspace learning [7], sparse coding [8], nonnegative matrix factorization [9], hashing [10], etc. In this paper, we will focus on the feature selection and subspace learning problem.
  — To handle the redundant and noisy features, **feature selection** is desired. Feature selection assigns different feature weights to different features, so that the useful features will be emphasized while the redundant and noisy features will be restrained [6].
  — To handle the the high-dimension problem of the feature vectors, the **subspace learning** could be employed for dimensionality reduction. Subspace learning maps the input feature vectors into a lower dimensional space, by using an optimal linear mapping matrix [7].

Many feature selection and subspace learning methods have been proposed to refine the original features, which could be classified into two types — supervised and unsupervised representation methods. The supervised method uses the class labels to guide the learning procedure, however, in database retrieval problems, the objects are usually not annotated, thus unsupervised representation is more suitable in this task.

- **Ranking Score Learning**: To compute the ranking score of a database object to a query, a distance or similarity measure could be employed to compare them, such as Euclidean distance, cosine similarity, etc. This type of methods is called pairwise similarity, and they only consider the query and objects to compare, while neglecting the manifold structure of the database. To handle this problem, the manifold ranking (MR) has been proposed by Zhou et al. [11], so that the ranking score could be learned with respect to the manifold structure of the database, which is characterized by a nearest neighbor graph constructed from the database. More over, Yang et al. [5] proposed the Local Regression and Global Alignment (LRGA) based ranking method to further improve the manifold ranking by using the local linear regression model for the ranking score learning problem.

The representation parameter is usually learned first, and then used to represent both the query and database objects. Based on the new representation, some ranking score learning algorithm will be applied for the ranking problem. Thus the representation and the ranking are conducted sequentially and independently. An important assumption behind this strategy is that the representation and the ranking are independent from each other, thus the possible inner relationships between them, which is not clear yet, has been ignored. It’s very interesting to notice that in [5], Yang et al. has applied the same LRGA model for both ranking and subspace learning. However, this model has been applied to the ranking and subspace learning respectively. In this paper, we argue that the representation and ranking should be considered in an unified way, so that we could investigate the possible relationships between them. Given a representation method, the ranking should be adjusted to the representation parameter. Moreover, given the ranking scores, the representation parameters should also be refined according to the ranking results.

To this end, we try to propose an unified framework for both the representation parameter learning and the ranking score learning, by constructing an unified objective function. The object representations parameterized by representation parameters will be used to compute the ground distances between query and database objects, and the the ground distances will be further used to regularize the ranking scores. At the same time, the ranking score will also be regularized by the manifold structure of the database. In this way, an unified objective function is built. The objective function will be optimized with regard to representation parameter and the ranking score alternately in an iterative algorithm. When the representation parameter is optimized, ranking score will be fixed, and then their role will be switched. Once the representation parameter is learned in the training procedure, it will be used to represent the new query object and rank the database objects. The contribution of this paper is listed as follows:

1. An unified framework for representation and ranking is proposed. Though we only discuss the feature selection and subspace learning as examples of representation, it could be extended to other representation methods easily, such as sparse coding, nonnegative matrix factorization, etc.
2. An iterative algorithm is proposed for the learning of representation parameters and ranking scores.

The remainder of this paper is organized as follows: In Section 2, we present the unified framework for representing and ranking. In Section 3, we apply the proposed framework to the brain tumor retrieval and nearest neighbor protein classification applications and show the experimental results. The conclusions and future works are given in Section 4.

## 2. Unified Framework for Representing and Ranking

In this section, we will introduce the novel framework for data ob ject representation and ranking in database retrieval and nearest neighbor classification tasks.

### 2.1. Objective Function

Suppose we have a database with *N* database objects, we denote it as 𝒟 = {**x**_1_,…, **x**_N_} ∈ ℝ^P^, where **x**_i_ = [*x*_*i*1_,…, *x*_*iP*_ ]^T^ ∈ ℝ^*P*^ is the P dimensional feature vector of the *i*-th database object. Given a query object, we denote it as **y** ∈ ℝ^*P*^, where **y** = [*y*_1_,…, *y*_*P*_]^T^ ∈ ℝ^*P*′^ is the *P* dimensional feature vector the query object. The task of database retrieval is to rank the database objects in *𝒟* according to the similarity between **y** and each **x**_*i*_ ∈ *𝒟*, and then return then few top ranked ones as retrieval results. To this end, we need to learn the nonnegative ranking score for each **x**_*i*_, denoted as *f*_*i*_, as the similarity measure between **y** and **x**_*i*_. The ranking scores of all the database objects are further organized as a ranking score vector 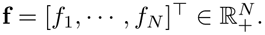 Moreover, instead of using the original features of query object **y**and the database object **x**_i_, we also consider to represent them by feature selection or subspace learning. The represented query and database objects are denoted as **y**^Θ^ ∈ ℝ^*P*′^ and **x**Θ ∈ ℝ^*P*′^, where Θ is the representation parameter, and *P*’ is the dimension of the feature space of the new representation.

To learn the representation parameter Θ and the ranking score vector **f** in an unified way, we will formulate the learning problem by an unified objective function. We will consider the following two regularization terms when constructing the objective function:

**Ground distance regularization**: Given a query object represented as **y**^Θ^, and a database object represented as 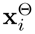, parameterized by Θ, we could compute the squared Euclidean distance between them as the ground distance: 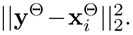. If the ground distance of query to *i*-th database object is short, it’s natural to expect the ranking score of i-th database objective is large; and vice versa. We model the regularization of ground distance with the following minimization problem:

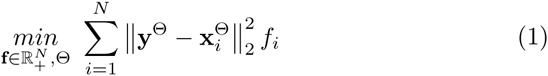

**Manifold regularization**: Based on the manifold assumption [12], which assumes that all the database objects lie on a low-dimensional manifold, we also try to regularize the ranking scores by manifold information. The manifold can be approximated linearly in a local area of the feature space of the database objects. Therefore, we assume that a database object x_*i*_ can be approximated by linearly reconstructing from its *K* nearest neighbors 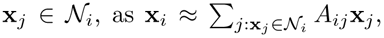, where *A*_*ij*_ is the reconstruction coefficient which summarizes the contribution of *x*_*j*_ to the reconstruction of X_*i*_. Following Locally Linear Reconstruction (LLR) [13], the coefficients *A*_*ij*_, *j* = 1,…, *N* could be obtained by minimizing the squared reconstruction error as:

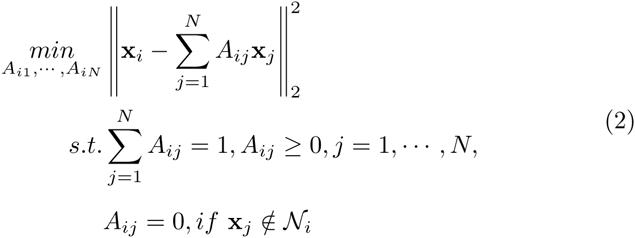

This problem could be solved as a Quadratic programming (QP) problem. The solved reconstruction coefficients are organized in a matrix 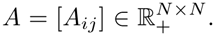. With the reconstruction coefficient matrix, we could formulate the manifold assumption to ranking scores by

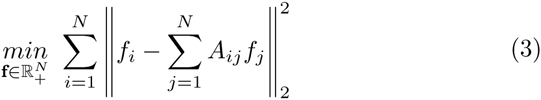

By solving this problem, we imply that a ranking score *f*_*i*_ could also be recontracted from the ranking scores *f*_*j*_ of its neighbors X_*j*_ ∈ *N*_*i*_. The manifold assumption is imposed to the ranking score by sharing the same local linear reconstruction coefficients *A*_*ij*_ between the feature space and the ranking score apace.

By combining the two regularization terms in (1) and (3), we could have the following objective function for the learning of f and Θ:

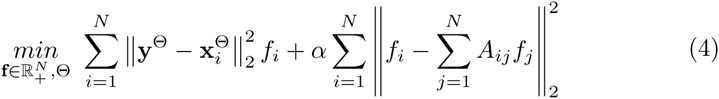

where α is a trade-off parameter.

We also suppose we have a query set with *M* query objects for the training procedure, denoted as *Q* = {**y**_1_,…, **y**_*M*_} ∈ ℝ^*P*^, where **y**_*k*_ = [*y*_*k*_, 1,…, *y*_*k*_, *P*]^T^ ∈ ℝ^P^ is the *P* dimensional feature vector of the *k*-th data object. When *k*-th query **y**_k_ is available in the training query set, we denote the ranking score vector for the *k*-th query object as 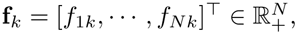, where *y*_*ik*_ is the ranking score of *i*-th database object against *k*-th query object. We define the ranking score matrix as 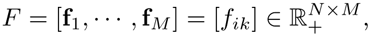, with its *k*-th collum as the ranking score vector of *k*-th query. Then the objective function could be extended to the following one by applying the objective function to each query and summing them up:

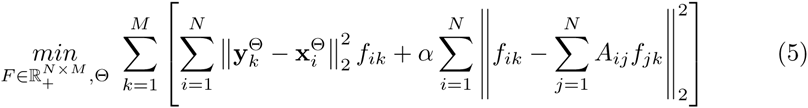

By minimizing the ob jective function in (5), we try to find the optimal ranking scores for the queries in **Q**, and the representation parameter Θ for both the query and databases objects in **Q** and 𝒟 simultaneously.

### 2.2. Optimization

To optimize the objective function (5), we adopt the alternate optimization strategy. **F** and Θ will be optimized alternatively in an iterative algorithm, and in each iteration, one of them will be solved or updated, while the other fixed, then their role will be switched.

#### 2.2.1. Optimizing F while fixing Θ

By fixing the representation parameter θ, and defining the ground distance matrix 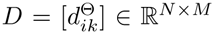, with 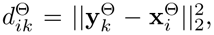, the problem (5) could be rewritten in matrix formula as,

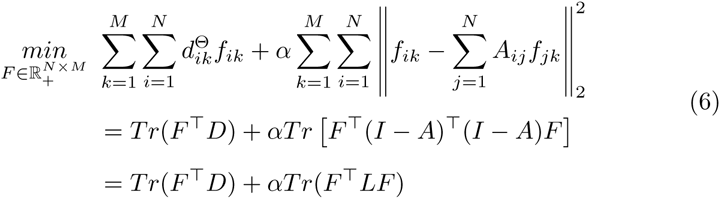

where *L* = *I*−2*A*+*A*^T^*A* ∈ R^*N*×*N*^. We introduce the lagrange multiplier matrix Φ = [*φ*_*ik*_] ∈ R^*N*×*N*^ for the constrain of 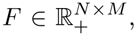, where *φ*_*ik*_ is the lagrange multiplier for constraint *f*_*ik*_ ≥ 0. The lagrange function *L* of the optimization problem is

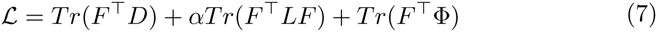

By setting the derivative of *L* with respect to *F* to zero, we have

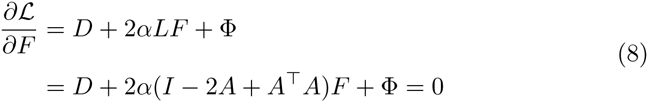

Using the KKT condition [Φ] ∘ [*F*] = 0, where [·] ∘ [·] denotes the element-wise matrix product, we get the following equation:

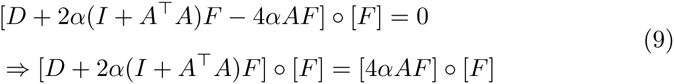

which leads to the following update rule for *F*

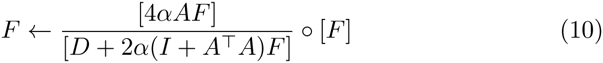

where 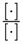 denotes the element-wise matrix division.

#### 2.2.2. Optimizing Θ while fixing F

To optimize Θ, we first need to specify the form of data representation which transfer a the original feature vector **X** ∈ ℝ^*P*^ to it newly represented feature vector **X**^Θ^ ∈ R^*P*′^, which is parameterized by Θ. Here we consider the feature selection and subspace learning as data representation methods, which are introduced as follows:

**Feature Selection**: Given a *P* dimensional feature vector **X** = [*x*_1_,…, *x*_*P*_]^T^ of a object, not all the features are relevant to the task in hand, and many of them might be noisy features. We try to assign each feature with different feature weight, so that the important features will be emphasized and the noisy features will be restrained. To this end, we introduce the nonnegative feature weight vector 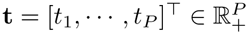 to parameterize the feature selection, where *t*_*p*_ is the weight for the *p*-th feature. The constrains *t*_*p*_ ≥ 0 and 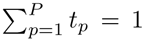 are introduced to **t**to prevent the negative weight. The feature vector could then be represented as

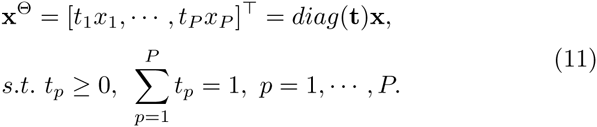

In this case, the representation parameter Θ is **t**. We apply the feature selection to both the query and the database objects, and then the ground distance between the *k*-th query object **y**_*k*_ and the *i*-th database object **X**_*i*_ will be computed as

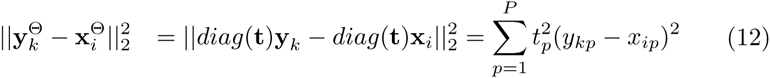

By replacing **t**by Θ, substituting (12) to (5), fixing *F* and removing the irrelevant term, (5) could be turned to the following optimization problem,

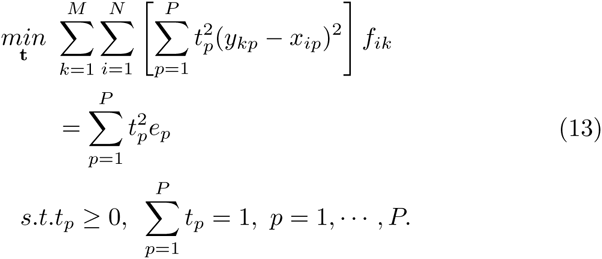

where 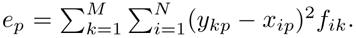
This problem could be efficiently solve as a standard QP problem as well.

**Subspace Learning**: Given the feature vector a data object **X** ∈ ℝ^*p*^, subspace learning [7] tries to map it into a *P*′’-dimension data space by a orthometric transformation matrix *W* ∈ ℝ^*P* x *P*′^ as

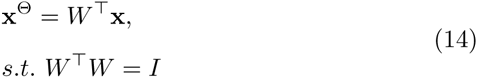

where *I* is an identity matrix of order *P*^’^. In this case, the representation parameter is *W*. By applying the subspace learning to both query and database objects, we have the ground distance between **y**_*k*_ and **x**_*i*_ defined as

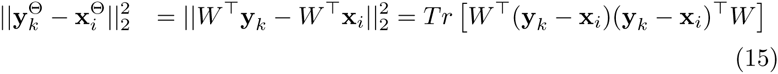

By replacing Θ by *W*, substituting (15) to (5), fixing *F*, and removing the term irrelevant to *W*, (5) could be turned to the following optimization problem,

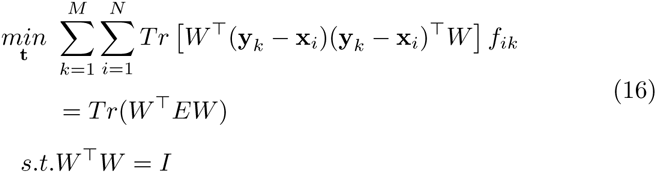

where 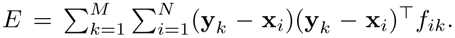
This problem could be obtained by solving the generalized eigenvalue decomposition problem,

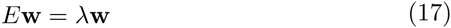

where λ is a eigenvalue and w ∈ ℝ^*p*^ is its corresponding eigenvector. Assume that the *P*′ smallest eigenvalues are ranked in a ascending order, as λ_1_,…, λ_*P*′_, and the corresponding eigenvectors are denoted as **w**_1_,…, **w**_*P*′_. Then the solution of (16) could be obtained as *W* = [**w**_1_,…, **w**_*P*′_] ∈ R^*P*x*P*′^.

### 2.3. Algorithm

Based on the optimization results, we could develop the iterative algorithm for the training procedure of unified object representation parameter Θ and the ranking score matrix *F*. The algorithm is summarized in Algorithm 1.

#### **Algorithm 1**UR^2^: off-line learning algorithm

**Input**: Database object set *𝒟* = {**x**_i_,…, **x**_N_}.

Input: Query object set *Q* = {y_*i*_,…, y_*M*_}.

Construct the nearest neighbor graph for *𝒟* and compute its reconstruction coefficient matrix *A*.

Initialize the ranking score matrix *F*^0^.

Initialize the representation parameter Θ^0^ and compute the initial ground distance matrix *𝒟*^0^.

for *t* = 1,…, *T* do

Update the ranking score matrix *F*^*t*^ based on the previous ground distance matrix *𝒟*^*t*–*t*^ and ranking score matrix *F*^*t*–1^, as in (10).

Update the representation parameter Θ^*t*^ by fixing *F*^*t*^, as in (13) or (17). Update the ground distance matrix *𝒟*^*t*^ based on the newly updated representation parameter Θ^*t*^.

**end for**

**Output**: The ranking score matrix *F*^*T*^, and the representation parameter Θ^*T*^.

### 2.4. Ranking new query object

We have introduced the off-line training procedure of Θ given a set of training query objects. In this subsection, we will discuss how to represent and rank a new query object y in the on-line retrieval procedure. In fact, we assume that the new arrived query won’t effect the representation parameter, and we use the parameter Θ learned using the training query objects to represent it as y^Θ^, based on feature selection or subspace learning. To learn its ranking score vector **f**, we simply solve the optimization problem in (4) while fixing Θ as learned by Algorithm 1. We define a ground distance vector for y^Θ^ against all the represented database objects as **d** = [*d*_1_,…, *d*_*N*_]^T^ ∈ ℝ^*N*^, where 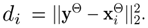. (4) then could be rewritten as

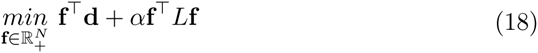

Its lagrange function *ℒ* of is

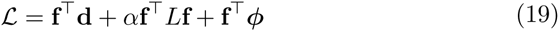

where ϕ ∈ ℝ^*N*^ is the lagrange multiplier vector for constrain f ≥ 0. By setting the derivative of *L* with respect to f to zero, we have

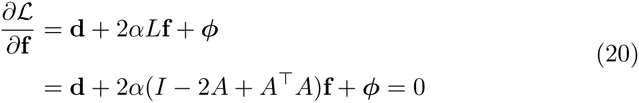

Using the KKT condition [ϕ]. [**f**] = 0, we get the following equation:

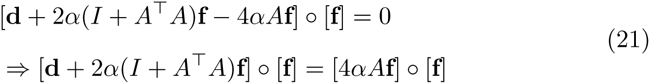

which leads to the following update rule for f

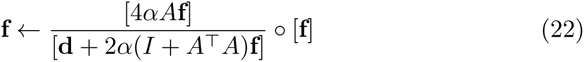

Based on the update rule, we could have the on-line ranking algorithm for query y, as summarized in Algorithm 2.

## 3. Experiments

In this experiment, we will evaluate the proposed methods for the brain tumor retrieval task and the nearest neighbor protein identification task.

### **Algorithm 2** UR^2^: on-line ranking algorithm

**Input**: Database object set *𝒟* = {x_i_,…, x_*N*_ } with its Laplacian matrix *L*.

**Input**: Query object **y**.

**Input**: The representation parameter Θ.

Initialize the ranking score vector f^0^.

Compute the ground distance vector d based on Θ.

for *t* = 1,…, *T* do

Update the ranking score vector f^*t*^ based on the ground distance vector d and previous ranking score vector f^*t*−1^ as in (22).

**end for**

**Output**: The ranking score vector f^*T*^.

### 3.1. Experiment I: Brain Tumor Retrieval

MRI has been one of the the most popular means for the diagnose of human brain tumors. However, the diagnosis of a brain tumor relies strongly on the experience of radiologists. In clinical practice, it would be significant helpful to have a retrieval system for brain tumors in MRI image which could return the tumors of the same pathological category as the query image. The doctors then can use the relevant MRI images returned by the retrieval system and the diagnosis information associated to these relevant images for the diagnosis for the current case [3]. In this experiment, we will evaluate the proposed method as MRI image representation and ranking method for the brain tumor retrieval system.

### 3.1.1. Dataset and Setup

Three types of brain tumors have been studied widely due to their high incidence rate in clinics, which are gliomas, meningiomas, and pituitary tumors. In this experiment, we use a dataset of 1014 MRI slices of the three types of brain tumors. There are 220 MRI slices of meningiomas, 475 MRI slices of gliomas, and 319 MRI slices of pituitary tumors in the dataset. The tumor regions in the images were manually outlined by drawing the the tumor boundaries. In this experiment, we define two tumor region as relevant if they contains tumors of the same type, otherwise, they are defined irrelevant. Given a query tumor region, the brain tumor retrieval task is to retrieve relevant tumor regions from the database. To this end, we extract visual features from the tumor region, including the following ones:

- **Intensity Features**: To extract the intensity features from the tumor region, we calculate the mean and variance of the normalized intensities of the tumor region pixels.
- **TeXture Features**: To extract the texture feature from the tumor region, we first calculate the Gray Level Co-occurrence Matrix (GLCM) and wavelet coefficients, and then some statistical parameters including mean, variance, entropy, correlation, etc, are estimated and used as texture features.
- **Shape Features**: To extract the shape features from the tumor region, we first calculate the shape signature from the points of the tumor boundary by using the radial distance, then perform the wavelet decomposition to the shape signature, and finally compute the mean and variance of the wavelet coefficients in each sub-band as shape features.
- **Bag-of-Words Features**: We also employ the bag-of-words model to extract the visual features from the tumor region. The key points are first detected, then the Scale-Invariant Feature Transform (SIFT) descriptor of each key points are calculated as “words”, and finally they are quantized to a dictionary and the quantization histogram is used as the bag-of-words feature.

All these features will be concatenated to obtain the visual feature vector of each brain tumor region in the MRI image. Using the proposed method, we perform the feature selection or subspace learning to the visual feature vector of query and database tumor regions to obtain the new representations, and learn the ranking scores of the database tumor regions according to the query tumor region for the ranking problem. Based on the ranking scores, the database tumor regions are ranked in a descending order of the ranking score, and the top few ones will be returned as relevant ones.

To conduct the experiment, we need a database, a training query set used to learn the representation, and a test query set to evaluate the retrieval performance. To this end, we randomly split the entire dataset into three subsets, one with 50% slices as database, one with with 25% slices as training query set, and another one with 25% slices as test query set. The database training query test qeury set split will be repeated randomly for ten times to reduce the bias of each split.

To evaluate the retrieval performances, we used the Receiver Operating Characteristic (ROC) and the recall-recision curves. The ROC curve is created by plotting True Positive Rates (TPR) against the False Positive Rates (FPR) of different numbers of returned tumors. The recall-precision curve is created by plotting precision against recall of different numbers of returned tumors. The TPR, FPR, precision and recall are defined as follows:

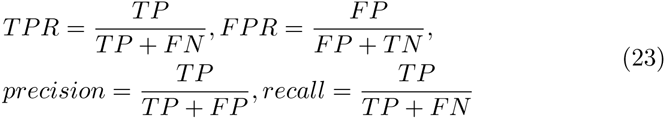

where *TP* is the number of returned tumors relevant to the query, *TN* is the number non-returned tumors irrelevant to the query, *FP* is the number of returned tumors irrelevant to the query, while *FN* is the non-returned tumors relevant to the query. Besides the curves, we also employ the Area Under the ROC Curve (AUC) and the Mean Average Precision (MAP) as the single measures for the retrieval task.

## 3.1.2. Results

In the experiments, we compare our unified framework for both representation and ranking of tumor region against several representation and ranking methods. The UR^2^ method with Feature Selection is donated as 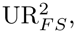, and UR^2^ method with Subspace Learning is donated as 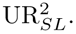. Since our methods are based on manifold learning of ranking score and representation parameters, we compare them against several manifold-based ranking and presentation methods, including:

- a feature selection method, Laplacian Score for Feature Selection (LSFS) [14],
- a subspace learning method, Locally Linear Embedding (LLE) [15],
- a ranking score learning method, LRGA [16], and
- the naive combinations of LRGA with LSFS and LLE respectively, denoted as “LRGA+LSFS” and “LRGA+LLE”.

show the results (average ROC and recall-precision curves) obtained by applying our methods 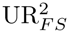 and 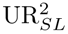 to the tumor region retrieval problem compared to other manifold-based representation and ranking score methods with tensity, texture, shape and bag-of-word histogram features. LLE has been chosen as a baseline since it has been extensively used in previous manifold learning works. Figure. 1 confirms the advantages of unfired representation and ranking approaches w.r.t. competing methods. For example, in the case of ROC our 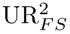 outperforms other methods consistently with different FPR values, which is followed by 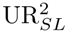. In the case of recall-precision curve, 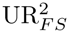 is more closer to the top right corner of the figure than any other methods. We should note that the proposed unified framework outperform not only the independent presentation and ranking methods (LRGA, LSFS and LLE), but also their naive combinations (LRGA+LSFS and LRGA+LLE). We explain this with the fact that our approaches, differently from other independent representation and ranking methods, take into account both representation and ranking problems simultaneously, so that the representation parameters and ranking scores could be learned optimally. Moreover, it is worth noting that the manifold ranking method (LRGA) outperforms the feature selection and subspace learning methods (LSFS and LLE) with pairwise distance as similarities, which highlights the importance of considering the manifold structure of the database when ranking. It’s also interesting to notice that for this task in hand, feature selection works better than subspace learning. The possible reason is that we have extracted many visual features from the tumor region while only few of them are relevant to the pathological type of the tumors. Similar conclusions can be made for the AUC and MAP values of the methods (see boxplots of AUC and MAP in Figure. 2). Also in this case the unified approaches of representation and ranking outperform independent representation and ranking methods.

**Figure 1:**
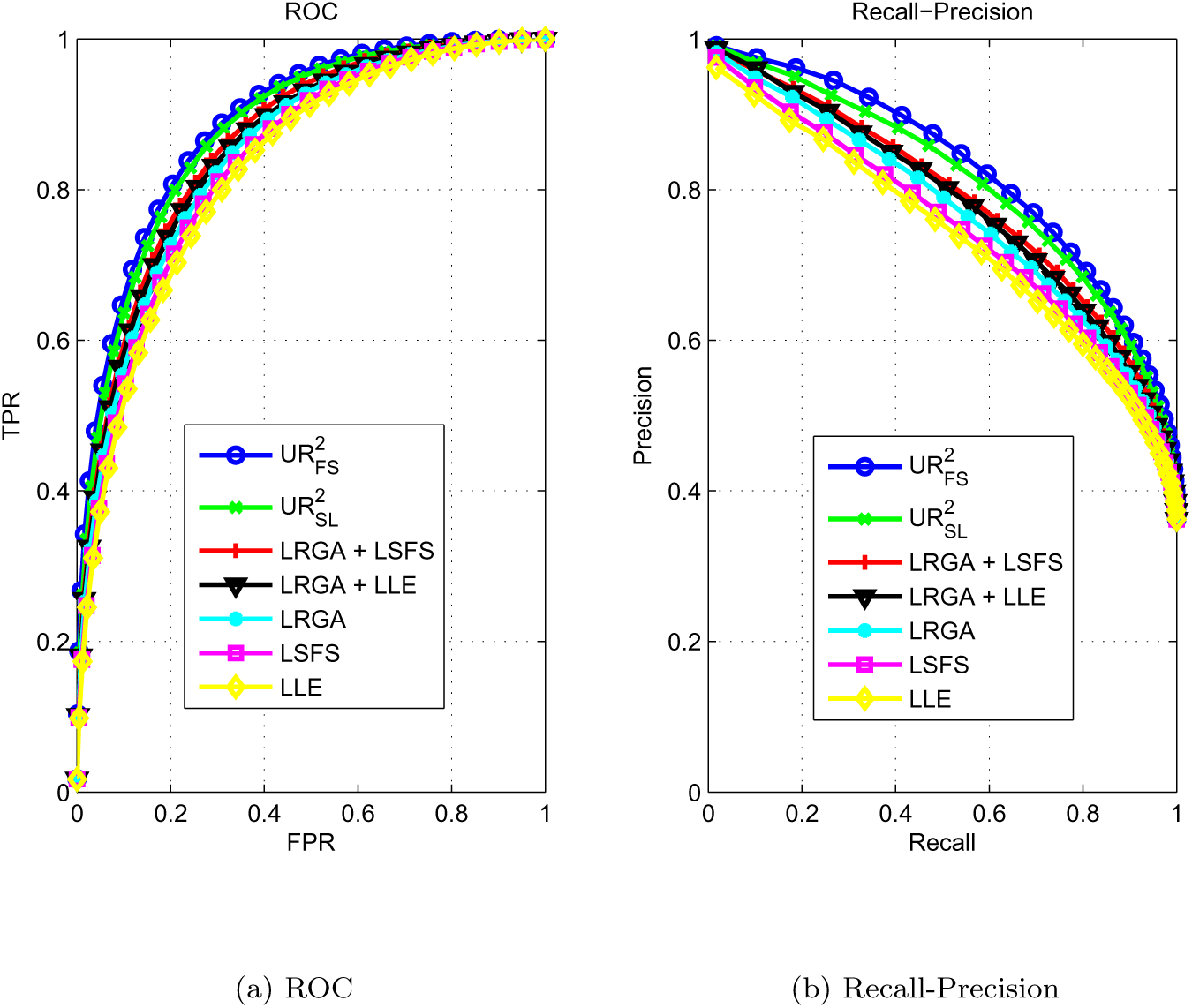
The ROC and recall-precision curves on brain tumor retrieval problem.

**Figure 2:**
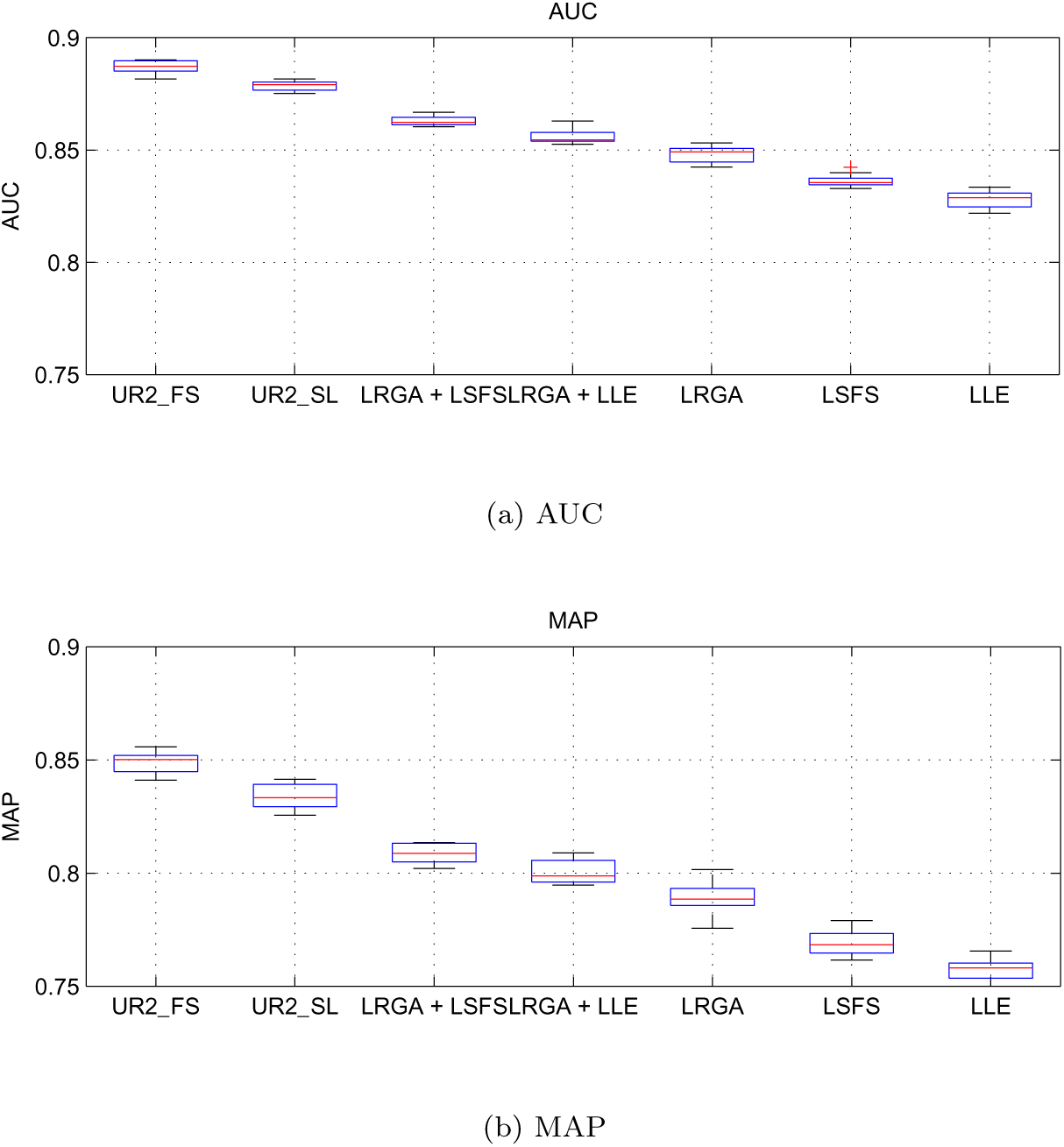
The AUC and MAP values on brain tumor retrieval problem.

## 3.2. Experiment II: Protein Identification

Identification the protein sample by using bio-sensor is very important for biochemical research and disease diagnose. In this experiment, we will evaluate the usage of proposed methods for the nearest neighbor classification based identification using the bio-sensor array data.

### 3.2.1. Dataset and setup

In this experiment, we collect a dataset of 100 protein samples, belonging to 9 different proteins. The 9 proteins are SubtilisinA (Sub), Fibrinogen (Fib), Hemoglobin (Hem), Cytochrome C (Cyt), Lysozyme (Lys), Horseradish perox idase (Hor), Bovine serum albumin (Bov), Lipase (Lip) and Casein (Cas). The sample number of each protein varies from 6 to 16. The distribution of sample number of different proteins is shown in Figure 3.

**Figure 3:**
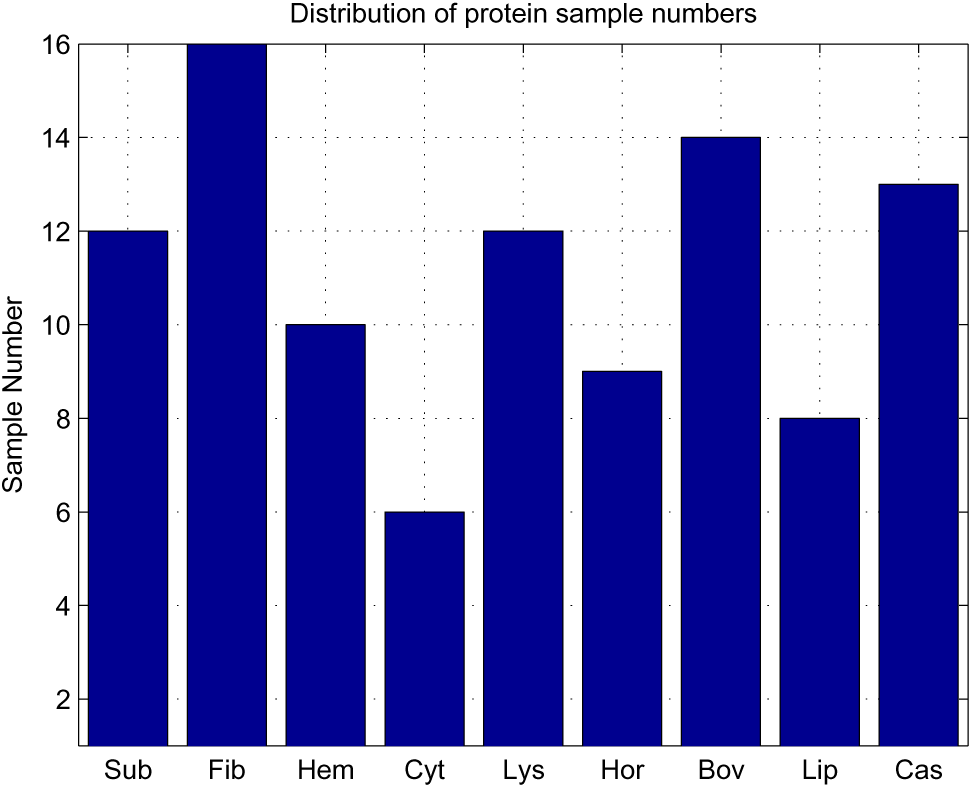
Distribution of protein sample numbers in the protein identification dataset.

Given an unknown sample, the task of protein identification is to classify the sample into one of the nine proteins in the training set. To this end, each sample will be tested against a bio-sensor array developed by Pei et al [17], called adaptive ensemble aptamers (ENSaptamers) which exploits the collective recognition abilities of a small set of rationally designed, nonspecific DNA sequences. The seven fluorescence intensities of a sample generated by seven ENSaptamers of the bio-sensor array are used as the original features and organized as a seven-dimensional feature vector. Then the feature vector of the query sample will be compared against all the feature vectors of the training samples in the datable and the most similar ones will be used for nearest neighbor classification.

To test the proposed methods, we employ the leave-one-out protocol to conduct the experiment. Each sample in the dataset will be used as a query sample in turns, while the remaining ones as training set. The training set will be further divided into training query set and database to learn the representation parameter. The training query set will contains 40% samples of the entire training set, while the database will contains 60% of the training samples. Once the representation parameter is learned by using the training set, it will be used to represent the query and the training samples. For the nearest neighbor classification of the query, the entire training set will be used as database. The ranking score of the database samples will be learned w.r.t the query, the ones with largest ranking scores will be returned and the query’s class label will be obtained by major voting of the returned samples.

The classification results are evaluated by the average classification accuracies of all the queries, which is defined as

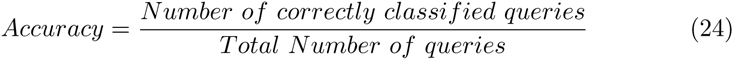

By varying the number of returned samples from the database, we could have different accuracies. The classification results will be reported using the curves of the accuracies against the returned sample numbers.

### Results

The accuracies of different methods with different different returned sample numbers are shown in Figure 4. It can be seen that both 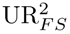 and 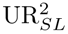 perform better than the best results of other methods at most cases, with 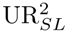 getting the overall best results. The combination of LLE/ LSFS and LRGA performs better than using individual representation or ranking methods, but could not beat the proposed unified framework. It indicates that using presentation and ranking methods together could boost the nearest neighbor classification performance, but the way to combine them is also very important. It’s also interesting to notice that 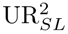 outperforms 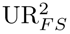 in this experiment, indicating that all the seven features of seven ENSaptamers are useful for the protein identification problem. This fact could also be verified by the fact that LLE outperforms LSFS. Moreover, it could be observed that when the returned sample number is small, the classifications are a kind of stable. However, when the re turned sample number is larger than 20, the classifications decrees significantly. This is because that for each query, there are at most 15 samples of the same protein in the database, which is defined as relevant to the query. When more than 15 samples are returned, the irrelevant samples will increase significantly and dominate the major voting of the nearest neighbor classification.

**Figure 4:**
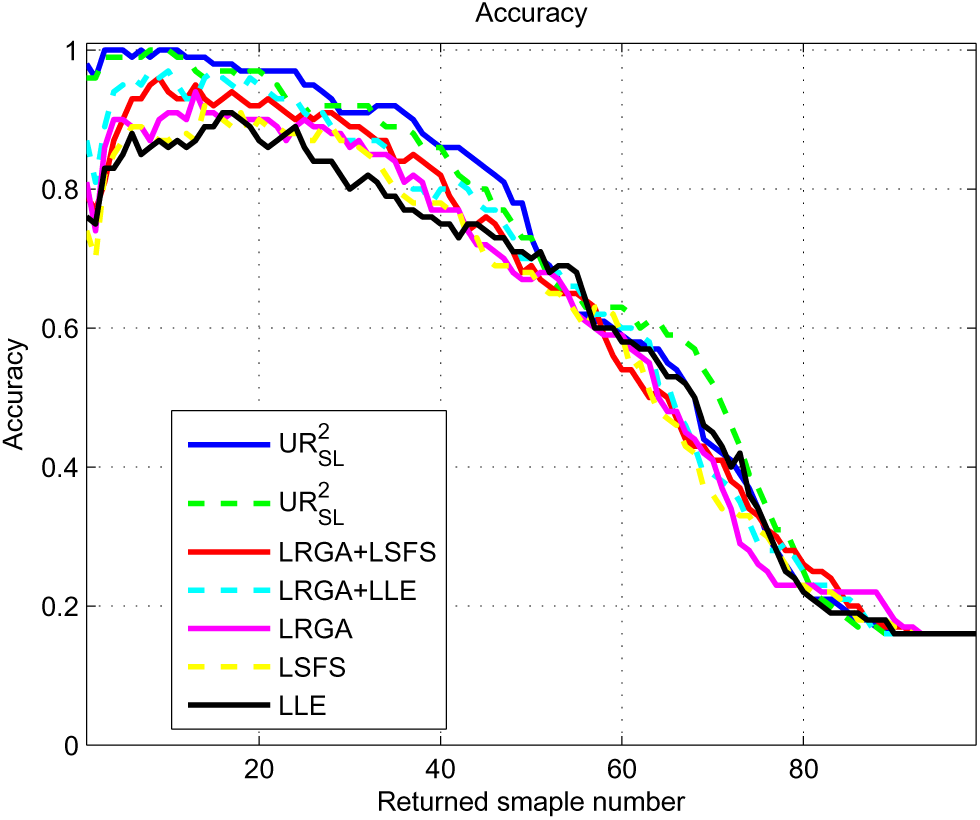
Curves of Accuracies against different returned sample numbers.

## 4. Conclusion and Future works

Representation learning and ranking score learning are two foundational problems for similar neighbor finding with many significant applications including database retrieval and nearest classification. Most research in the machine learning community have been focussed on the learning of representation parameters and ranking score respectively, which ignores the possible relationships between these two issues at all. In this paper, for the first time, we propose the unified framework for representation and ranking objects in database retrieval and nearest classification problems. It is shown in this work that using the proposed unified framework to learn the representation and raking parameters works well in this scenario. A significant advantage of the proposed method, as compared to methods to represent and rank objects respectively, is that, with different representation parameter to define the ground distance, the optimal ranking scores could be learned according to the representation parameter. Moreover, the representation parameter could also be adjusted according to the ranking scores.

For the future works, we would consider using sparse coding as the representation method instead of features selection and subspace learning, which is the stat-of-the-art representation method. Moreover, the optimization of the ranking score could possibly has close form, which is another direction desired to explore.

